# Heart rate variability as a candidate correlate of susceptibility to ASMR and music-induced frisson: an exploratory pilot study

**DOI:** 10.64898/2026.04.01.715955

**Authors:** L Amthor, O Brüngger, M Bühler, A Monn, B Provaznikova, G Kronenberg, S Olbrich, T Welt

## Abstract

**Background:** Autonomous sensory meridian response (ASMR) and music-induced frisson are sensory-affective phenomena characterized by tingling, chills, and pronounced emotional responses. Previous research has mainly focused on physiological changes during these experiences, whereas much less is known about whether baseline physiological state is associated with subsequent susceptibility.

**Objective:** To examine whether baseline autonomic flexibility, indexed primarily by heart rate variability (HRV), is associated with later ASMR/frisson responsiveness. Resting EEG measures were included as secondary exploratory markers.

**Methods:** Fifteen participants were recruited by convenience sampling; after artifact-based exclusion, 10 participants were included in the analyses. A 5-minute resting baseline EEG and ECG was recorded prior to stimulus presentation. Participants were then exposed to auditory and audiovisual ASMR stimuli, classical music excerpts, and a control stimulus, and reported whether they had experienced ASMR-typical sensations or frisson. Main analyses examined associations between baseline physiological parameters and a combined response-positive outcome. Exploratory analyses included participant-level correlations, comparisons between susceptible and non-susceptible participants, and stimulus-specific effect sizes.

**Results:** HRV-related measures showed the clearest and most consistent pattern of association with responsiveness. Higher baseline total HRV power was associated with a greater number of response-positive stimuli (r = 0.756, p = 0.011), with similar positive associations for high-frequency HRV (HF; r = 0.672, p = 0.033) and baseline heart rate slope (r = 0.751, p = 0.012). Stimulus-specific analyses likewise showed the most consistent positive baseline effects for total HRV power, with HF and heart rate slope pointing in the same direction. Frontal alpha asymmetry (FAA) was negatively associated with responsiveness (ρ = -0.862, p = 0.001), but EEG findings overall were less consistent than the HRV-related pattern and are best interpreted as secondary exploratory observations.

**Conclusions:** In this exploratory pilot sample, baseline HRV, particularly total HRV power, showed the most coherent physiological association with susceptibility to ASMR and music-induced frisson. The findings are consistent with the possibility that these experiences depend not only on stimulus properties, but also on pre-existing physiological state. Given the small sample and exploratory design, the results should be interpreted as hypothesis-generating and require replication in larger confirmatory studies.

## Introduction

Music has long been recognized as a powerful modulator of human affective and physiological states. Since the 1970s, research has shown that music can influence mood, reduce anxiety, and affect cognitive performance ^1,2^. Beyond these psychological effects, music also produces measurable autonomic changes. For example, listening to slow classical music has been associated with reductions in heart rate and blood pressure ^3^, and meta-analytic evidence supports cardiovascular effects of music-based interventions ^4^. Together, these findings indicate that music engages not only subjective emotional experience but also physiological regulation.

One particularly salient music-related phenomenon is frisson, a brief sensation of chills or shivers, often accompanied by piloerection, that typically occurs during emotionally intense moments in music ^5–7^. Frisson has been linked to specific musical features, including dynamic changes, the introduction of new instrumental or vocal elements, unexpected harmonic transitions, and marked melodic contrasts ^8,9^. Such features are thought to elicit strong emotional reactions through mechanisms of tension, expectation, and expectancy violation ^8,9^. Empirically, frisson has been associated with increased autonomic arousal, including elevations in heart rate and skin conductance, particularly when accompanied by objectively measurable piloerection ^10–13^.

A related but distinct phenomenon is autonomous sensory meridian response (ASMR), a pleasant tingling sensation that typically begins on the scalp and spreads across the neck and spine, often accompanied by relaxation and positive affect ^14–17^. Common triggers include whispering, soft repetitive sounds, and cues of personal attention such as simulated caregiving interactions ^14,16–19^. Although ASMR and frisson differ in duration, affective tone, and trigger profile, both involve a distinctive bodily-affective response characterized by tingling, chills, or shiver-like sensations ^14,20^. Frisson is usually brief and linked to emotional peaks, whereas ASMR is typically more sustained, calming, and immersive ^14,20^.

Research on both phenomena has focused primarily on what happens during the experience itself. In ASMR, studies have reported reduced heart rate alongside increased skin conductance, suggesting concurrent parasympathetic and sympathetic engagement ^15,17^. EEG studies have further identified alterations in alpha, beta, and gamma activity, consistent with a relaxed but attentive state ^21,22^. In frisson, neuroimaging studies have implicated reward-related regions, including the nucleus accumbens, and have demonstrated dopaminergic involvement during intensely pleasurable musical moments ^10,13^. Taken together, these findings suggest that ASMR and frisson recruit overlapping affective, autonomic, and reward-related mechanisms once the experience is already unfolding.

By contrast, far less is known about whether individuals differ in their physiological predisposition to experience ASMR or frisson in the first place. Most previous studies have classified individuals as ASMR-sensitive or non-sensitive, or as more or less prone to frisson, primarily on the basis of self-report (^16,18,19,23^). While this supports the existence of stable interindividual differences, it does not clarify whether such differences are reflected in measurable baseline physiological characteristics prior to stimulus exposure.

A particularly plausible candidate marker is heart rate variability (HRV). HRV is widely regarded as an index of autonomic flexibility and has been linked to emotional regulation, adaptive responsiveness, and the capacity to shift efficiently between physiological states. If susceptibility to ASMR and frisson depends not only on stimulus properties but also on an individual’s regulatory readiness, then resting HRV may capture an important component of this predisposition. Resting EEG measures may provide additional information about trait-like differences in cortical state or attentional style, but in the present context they are considered secondary exploratory markers.

The present exploratory study addresses this gap by examining whether baseline ECG/HRV measures and resting EEG parameters, recorded prior to stimulus presentation, are associated with the occurrence and frequency of subsequent ASMR/frisson experiences. Our central working hypothesis was that individuals with higher baseline autonomic flexibility, indexed primarily by HRV, would show increased susceptibility to ASMR/frisson. EEG measures, including frontal alpha asymmetry and spectral parameters, were included as secondary exploratory markers. By extending the focus from stimulus-evoked responses to physiological predisposition, this study aims to provide an initial mechanistic framework for understanding why some individuals appear more responsive than others.

## Methods

### Participants

Fifteen healthy volunteers were enrolled in the study. After data-quality control, five participants were excluded because of pronounced artifacts in the physiological recordings, leaving a final analytic sample of 10 participants (6 male, 4 female; *M* = 33.2 years, *SD* = 16.0). Age distribution was bimodal, comprising a younger subgroup (*n* = 7, *M* = 23.3 years, *SD* = 1.0) and an older subgroup (*n* = 3, *M* = 56.3 years, *SD* = 1.53).

### Stimuli and procedure

The experimental paradigm combined ASMR and music stimuli to probe individual susceptibility to ASMR/frisson-like experiences across a range of affective auditory contexts. Because the present study focused on baseline physiological predisposition, stimulus-specific characteristics were not analyzed in detail.

ASMR stimuli consisted of selected audio excerpts from publicly available ASMR recordings and were classified according to their dominant sensory and contextual characteristics. Ten stimuli representing common ASMR trigger profiles were selected. Five ASMR stimuli were presented in auditory-only form and five audiovisually.

The music set consisted of four classical excerpts varying in tempo, tonality, rhythm, and emotional character: Wolfgang Amadeus Mozart’s *Piano Concerto No. 21 in C major, KV 467* (2nd movement, Andante), Max Richter’s *Recomposed: Summer 3*, Léo Delibes’ “Flower Duet” from *Lakmé*, and Alessandro Marcello’s *Oboe Concerto in D minor* (Adagio, arranged for mandolin). Two music excerpts were presented in audio-only form and two with visual/contextual framing. A train sound served as a control stimulus to assess recognition and response specificity.

At the beginning of the experiment, participants completed a brief background questionnaire including music preferences and prior ASMR experience; these variables were not included in the present analyses. A 5-minute resting baseline was then recorded without stimulation while EEG and ECG data were acquired. Subsequently, stimuli were presented in two blocks: an auditory block, in which stimuli were delivered in randomized order via in-ear headphones, and an audiovisual block, in which stimuli were presented on a 13-inch screen at a viewing distance of approximately 1 m using the same headphones.

After completion of the auditory block, participants reported whether they had experienced ASMR during the auditory ASMR stimuli as a whole, whether frisson had occurred during each of the two classical music excerpts, whether the control stimulus had been correctly identified, and whether ASMR had occurred during the control stimulus. After each audiovisual stimulus, participants reported whether they had experienced ASMR-typical tingling sensations or frisson.

### Physiological recording and preprocessing

Physiological recordings were obtained between 09:00 and 18:00. During baseline acquisition, participants sat in a reclined position in a darkened room with their eyes closed and were instructed to remain as still as possible. Room temperature was maintained between 20 and 23°C.

EEG was recorded using an eego amplifier system (ANT Neuro, Hengelo, Netherlands) with 64 Ag/AgCl electrodes, including vertical electrooculography (EOG). Electrode placement followed the 10–20 system, with CPz as reference. Horizontal eye movements were recorded using two bipolar electrodes placed near the outer canthi. ECG was recorded simultaneously using two bipolar electrodes placed on the wrists. Data were acquired at a sampling rate of 4 kHz, and electrode impedances were kept below 50 kΩ.

Resting EEG data were preprocessed in DeepPSY software (Zollikerberg, Switzerland, 2024). Processing included low-pass filtering at 70 Hz, high-pass filtering at 0.5 Hz, and a 50 Hz notch filter. Data were segmented into 1-second epochs and visually inspected for movement artifacts; up to 10% of segments were removed where necessary. Independent component analysis (ICA) was then performed after resampling to 250 Hz, and artifact-related components were identified based on scalp topography and comparison with the raw signal. Typically, 2–3 components reflecting eye movements or muscle activity were removed. EEG-derived variables were treated as secondary exploratory markers.

Baseline ECG/HRV measures constituted the primary physiological markers of interest. ECG signals were analyzed in DeepPSY software following automated R-peak detection, with visual inspection and manual correction where necessary. The extracted parameters were heart rate (HR; beats per minute), rate of change in heart rate (heart rate slope), low-frequency power (LF; 0.04–0.15 Hz), high-frequency power (HF; 0.15–0.4 Hz), normalized low-frequency power (LFnu = LF / [LF + HF] × 100), and total HRV power.

### Outcome definition

Participants were classified as response-positive when they reported ASMR-typical tingling sensations and/or frisson following stimulus exposure. For the primary participant-level analyses, responsiveness was operationalized as the number of stimuli associated with a response-positive report. ASMR and frisson were combined pragmatically for the main analyses because of their partial phenomenological overlap, the pilot character of the study, and the limited sample size.

### Statistical analysis

Statistical analyses were conducted using R (version 4.5.1; R Core Team, 2025) and RStudio (version 2025.09.0; Posit Software, PBC). All analyses were exploratory and used a two-sided significance threshold of α = 0.05.

The primary analytical focus was on associations between baseline ECG/HRV measures and subsequent ASMR/frisson reports. Resting EEG measures were analyzed secondarily and interpreted as exploratory. Because of their partial phenomenological overlap, the pilot character of the study, and the limited sample size, ASMR and frisson were combined into a single response-positive outcome in the main analyses. This decision was intended to preserve sensitivity for detecting potential baseline physiological signals, but the combined endpoint should be interpreted cautiously because the two phenomena are not identical.

At the stimulus level, participants were classified according to whether a given stimulus was followed by a reported ASMR/frisson experience (“yes”) or not (“no”); for the auditory ASMR set, classification was based on a single combined block-level report. Stimuli with only a single response-positive event were excluded from this comparison. Group differences in baseline parameters were examined using independent-samples *t*-tests. Homogeneity of variance was assessed using F-tests, and Welch correction was applied where appropriate. Effect sizes were quantified using Hedges’ *g*, with positive values indicating higher baseline values in the response-positive group and negative values indicating lower baseline values in that group.

To capture more aggregated susceptibility, associations between baseline physiological variables and the number of stimuli eliciting ASMR/frisson were examined using Pearson and Spearman correlations. In addition, participants who never reported ASMR/frisson were compared with those who reported it at least once using the same statistical framework.

Given the exploratory and pilot nature of the study, no correction for multiple comparisons was applied. Results are therefore interpreted as hypothesis-generating, with a primary focus on effect sizes as indicators of potential signal strength rather than precise population estimates.

### Ethics

This study was conducted in accordance with the Declaration of Helsinki. No formal ethical approval had been obtained prior to data collection in 2024. Following a jurisdictional inquiry (Req-2025-01671), an Advisory Opinion was obtained from the Cantonal Ethics Committee Zurich (KEK; BASEC AO_2026-00015). In its opinion dated March 20, 2026, the committee stated that the project, as described in the submitted documents, was in principle consistent with the regulatory and ethical standards of the Swiss Human Research Act.

## Results

### Associations Between Baseline HRV and ASMR Susceptibility

Baseline cardiac autonomic indices were positively associated with ASMR/frisson responsiveness. Across participants, higher baseline heart rate variability (HRV) was associated with a greater number of stimuli eliciting ASMR/frisson responses (Figure 1). Total HRV power showed a strong positive correlation with responsiveness (*r* = 0.756, *p* = 0.011; Figure 1A). A similar pattern was observed for high-frequency HRV (HF; *r* = 0.672, *p* = 0.033; Figure 1B) and heart rate slope (*r* = 0.751, *p* = 0.012; Figure 1C). Taken together, these findings indicate that participants with higher baseline cardiac autonomic variability tended to respond positively to a broader range of ASMR/frisson-inducing stimuli.

**Figure 1.**
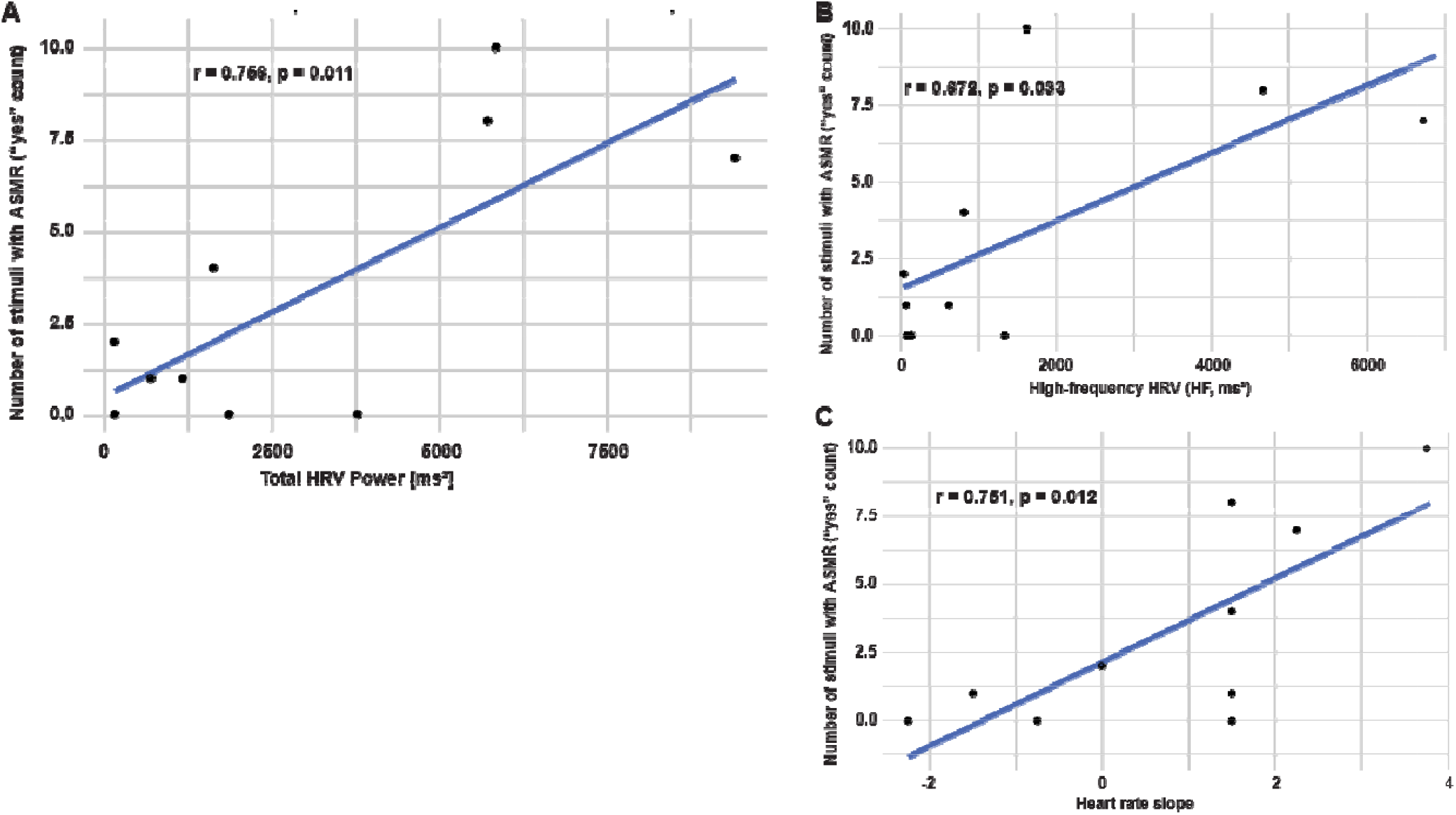
Association between baseline heart rate variability (HRV) parameters and susceptibility to ASMR/frisson experiences. **A.** Total HRV power showed a strong positive association with the number of stimuli eliciting ASMR/frisson responses. **B.** High-frequency HRV (HF) and **C.** heart rate slope showed similar positive trends. Lines represent linear regression fits with 95% confidence intervals Pearson correlation coefficients are reported in the text.

### Associations Between Baseline FAA and ASMR Susceptibility

Baseline frontal alpha asymmetry showed an inverse association with responsiveness. As a secondary cortical marker, frontal alpha asymmetry (FAA) was strongly negatively associated with the number of stimuli eliciting ASMR/frisson responses (Figure 2). Spearman rank correlation revealed a robust inverse relationship between baseline FAA and responsiveness (ρ = -0.86, p = 0.001, n = 10), indicating that lower baseline FAA values were associated with greater ASMR/frisson susceptibility. Thus, although autonomic measures showed the most consistent pattern across analyses, FAA emerged as the most notable exploratory cortical finding in the present sample.

**Figure 2.**
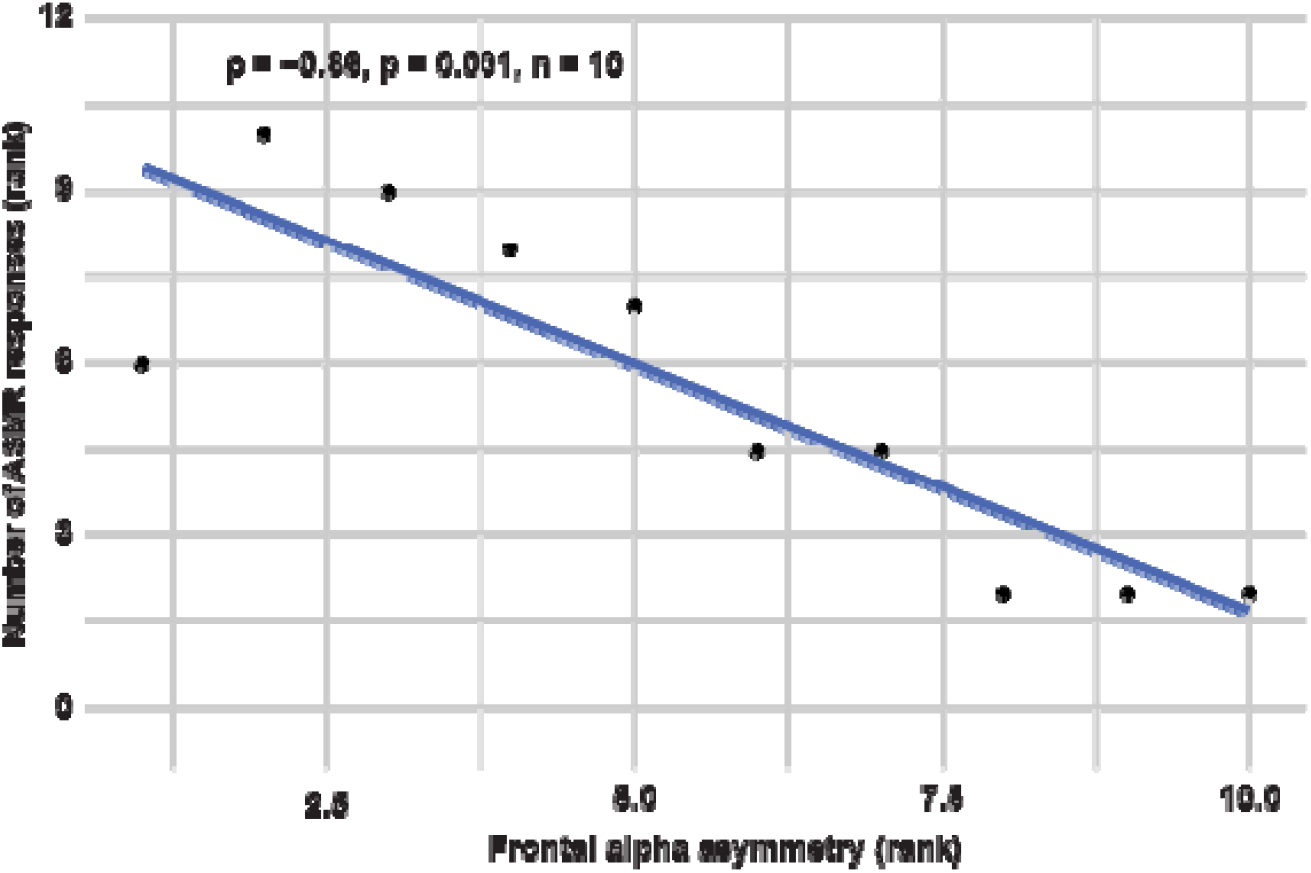
Negative association between frontal alpha asymmetry (FAA) and susceptibility to ASMR/frisson experiences. Spearman rank correlation revealed a strong Inverse relationship between baseline FAA and the number of stimuli eliciting ASMR/frisson responses (p = −0.88, p = 0.001). Shaded area represents the 95% confidence Interval.

### Baseline Differences Between Susceptible and Non-susceptible Participants

To evaluate systematic baseline differences, participants were classified based on their overall responsiveness (reporting at least one ASMR or frisson experience). As shown in Figure 3, susceptible participants exhibited a distinct physiological profile, with cardiac autonomic markers showing a consistent directional pattern. Specifically, susceptible individuals exhibited higher baseline values for heart rate slope (g = 0.94), high-frequency HRV (HF; g = 0.62), and Total HRV power (g = 0.45). These autonomic differences were accompanied by a large group effect in frontal alpha asymmetry (FAA; g = -1.61), with susceptible participants showing lower baseline values. Overall, these findings suggest that individuals susceptible to ASMR and frisson exhibit a specific baseline configuration of both autonomic and cortical parameters.

**Figure 3.**
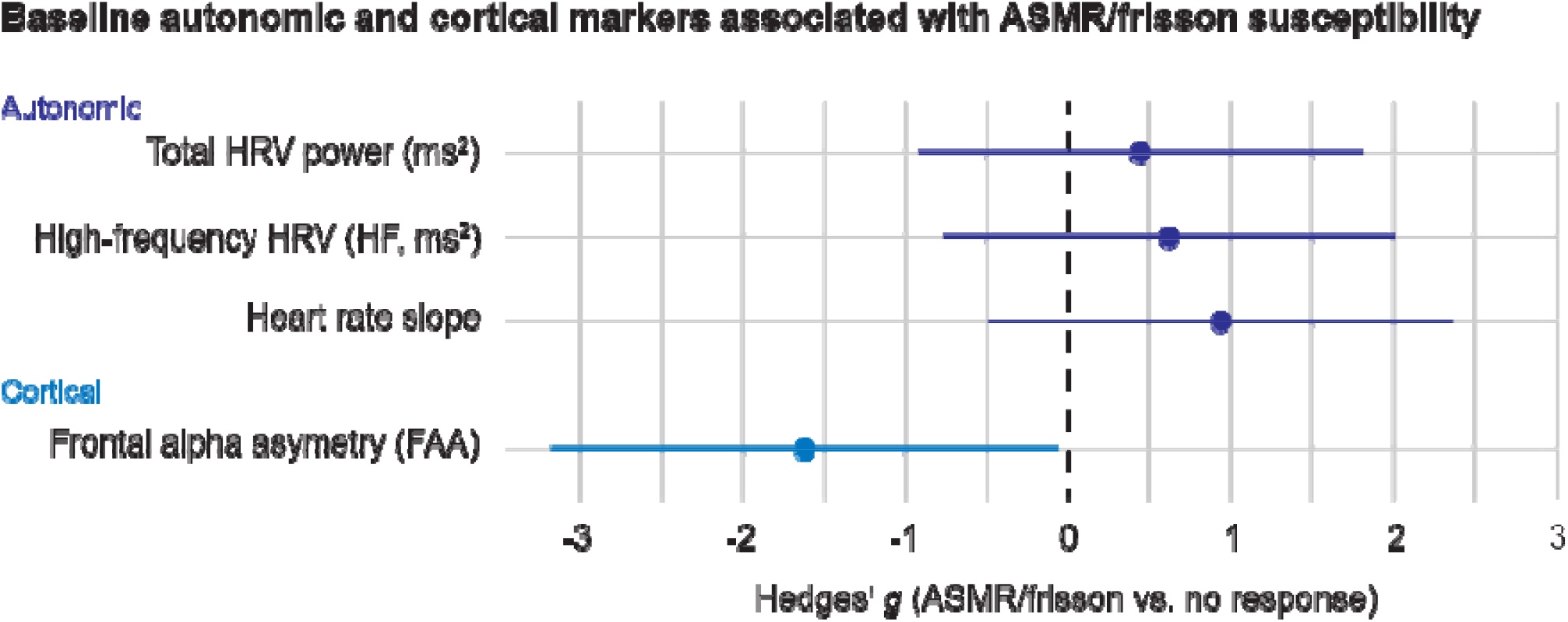
Effect sizes (Hedges’ *g*) for baseline differences between response-positive (ASMR/frisson) and response-negative participants, Positive values indicate higher baseline values preceding reported ABMR/-frisson experiences. HRV-related measures (total HRV power, HF, and heart rate slope) showed consistently positive effects, whereas frontal alpha asymmetry (FAA) showed a negative effect. Error bars represent 95% confidence intervals.

### Stimulus-Specific Baseline Effects

To evaluate the consistency of baseline physiological signatures across different auditory and audiovisual contexts, stimulus-specific effect sizes (Hedges’ g) were calculated for all measured parameters (Figure S1). Stimuli with only a single response-positive event were excluded from this analysis.

The cardiac autonomic signal remained remarkably stable across diverse trigger categories. Specifically, Total HRV power exhibited pronounced and consistent positive baseline effects (mean g = 1.89), with the largest positive effects observed for Audio ASMR, Eating sounds, and Classical music – spring (all p < 0.001). High-frequency (HF) power showed a similar pattern (mean g = 1.64); however, these effects did not reach significance after Welch correction for unequal variances. Additionally, heart rate slope showed consistently positive effects (mean g = 1.22), most notably for the Soft tactile sounds condition (p = 0.023) and the three aforementioned stimuli (p = 0.039).

In contrast, EEG-derived and non-HRV measures showed higher stimulus-dependency and predominantly negative effects. LFnu displayed a mean effect of g = -0.93, which was most pronounced for the Brushing stimulus (p = 0.046). Similarly, frontal alpha asymmetry (FAA) showed a significant negative baseline effect preceding ASMR reports during the Whispering condition (g = -0.59, p = 0.036), while other qEEG parameters displayed more varied patterns across the stimulus set.

**Supplementary Figure S1.**
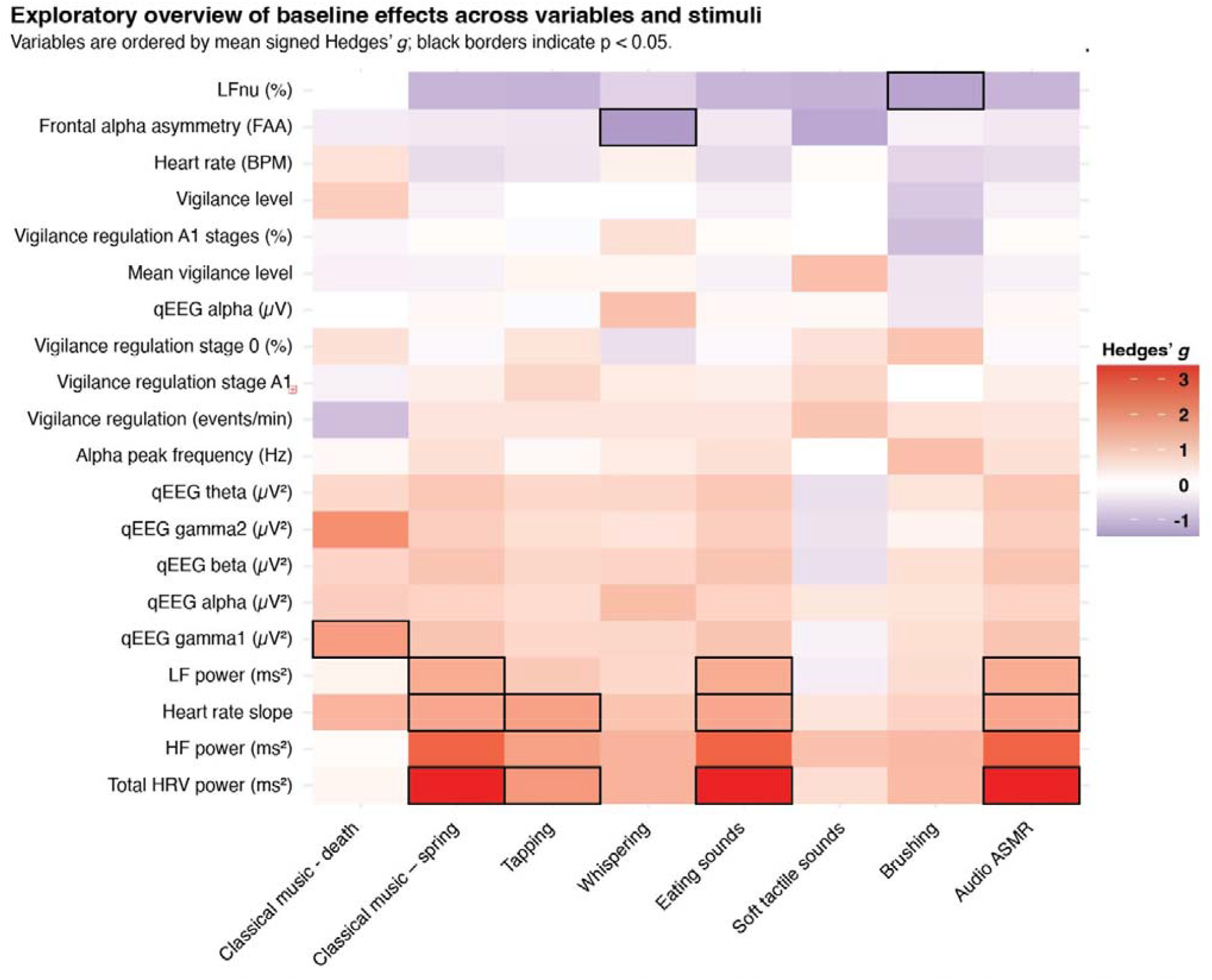
Exploratory overview of baseline effects across variables and stimuli Heatmap showing effect sizes (Hedges’ *g*) for baseline differences between response-positive (ASMR/frisson) and response-negative trials across all measured variables and stimulus conditions. Positive values indicate higher baseline values preceding reported ASMR/frisson experiences, whereas negative values indicate lower baseline values. Variables are ordered by mean signed Hedges’ *g* Black borders indicate statistically significant effects (p < 0.05).

## Discussion

The present exploratory pilot study examined whether physiological state prior to stimulus exposure is associated with subsequent susceptibility to ASMR and music-induced frisson. The main finding was that HRV-related baseline measures showed the most consistent pattern of association with later response-positive experiences. Across analyses, total HRV power, high-frequency HRV (HF), and baseline heart rate slope were positively related to responsiveness, whereas EEG findings were less consistent overall. Among all parameters examined, total HRV power emerged as the most coherent signal in this pilot sample, suggesting that baseline autonomic variability may represent a relevant physiological correlate of ASMR/frisson susceptibility. This finding adds a new perspective to the existing literature. Previous studies on ASMR have primarily focused on physiological changes during the experience itself, including reduced heart rate and increased skin conductance ^15,17^, while work on music-induced frisson has emphasized stimulus-evoked autonomic and reward-related responses during emotionally intense moments ^10–13^. In contrast, the present results suggest that baseline physiology may also matter. This extends the focus from response expression to possible response predisposition and is consistent with the idea that ASMR and frisson depend not only on the eliciting stimulus, but also on the physiological state in which that stimulus is encountered.

The prominence of HRV in the present dataset is physiologically plausible. HRV is widely used as an index of autonomic regulation ^24^, and higher resting HRV has repeatedly been associated with better emotion regulation, adaptive stress responding, and greater psychological resilience^25–30^. HF is commonly interpreted as reflecting predominantly vagally mediated influences ^24,31^, whereas LF-related metrics are more difficult to interpret and should not be treated as direct indices of sympathetic activity alone ^32^. In this context, the present pattern is compatible with the interpretation that individuals with greater baseline autonomic flexibility may be more likely to enter the immersive and affectively engaged state in which ASMR or frisson becomes reportable. Given the exploratory design and small sample, however, this interpretation should remain cautious and non-causal.

A notable aspect of the results is that the HRV signal was not confined to a single trigger type. Total HRV power showed pronounced positive baseline effects across several auditory and audiovisual categories, including Audio ASMR, Eating sounds, and Classical music – spring, and HF as well as heart rate slope pointed in the same general direction. This suggests that baseline autonomic variability may reflect a broader susceptibility factor rather than a narrow response tendency tied to one specific class of stimuli. At the same time, ASMR and music-induced frisson are not identical phenomena ^5–8,8,9,14^. Because response-positive events were aggregated across both, the present findings are best interpreted as indicating susceptibility to a shared class of sensory-affective experiences, without implying that both phenomena necessarily rely on identical mechanisms.

Resting EEG measures played a secondary role in the present study. Frontal alpha asymmetry (FAA) was inversely associated with responsiveness and represented the most notable exploratory EEG finding in the dataset. However, the EEG pattern overall was less consistent than the autonomic one: FAA showed a stimulus-specific effect in the Whispering condition, whereas other qEEG measures varied across analyses and stimuli. Prior work has linked FAA to aspects of affective style, attentional orientation, and related trait-like characteristics ^19,23,33,34^. In the present context, FAA may reflect an additional aspect of resting-state predisposition, although this interpretation remains exploratory and requires replication. In the present study, FAA is best understood as a secondary exploratory observation, whereas HRV constitutes the primary physiological signal.

Several limitations should be considered. First, this was a small exploratory pilot study based on a convenience sample, and only 10 participants were retained for statistical analyses after artifact exclusion. This limits statistical power, precision, and generalizability, and the proportion of exclusions may also have introduced selection bias. Second, the analyses were explicitly exploratory and covered multiple physiological parameters and stimulus categories, so the results should be interpreted as hypothesis-generating rather than confirmatory. Third, ASMR and frisson were partially combined into a single response-positive category, which was pragmatic in this pilot context but remains a conceptual limitation. Fourth, susceptibility was assessed by self-report, with most outcomes recorded after individual stimuli but the auditory ASMR set assessed at block level, and these reports may have been influenced by individual reporting thresholds and differences in the interpretation of bodily sensations. Finally, the cross-sectional design does not allow causal inference.

Future studies should therefore focus first on replicating the HRV findings in larger, preregistered samples with prespecified primary outcomes. Based on the present results, total HRV power appears to be the most promising primary candidate marker, with HF and baseline heart rate slope as relevant secondary autonomic measures. It will also be important to separate ASMR and music-induced frisson analytically in order to test whether the same baseline autonomic profile predicts both phenomena. More broadly, experimental studies using breathing-based modulation or HRV biofeedback may help clarify whether HRV is merely associated with susceptibility or contributes more directly to it ^33–36^.

In conclusion, this exploratory pilot study provides preliminary evidence that baseline heart rate variability is associated with susceptibility to ASMR and music-induced frisson. Among all measures examined, HRV-related indices—especially total HRV power—showed the most consistent relationship with later response-positive experiences. EEG findings, particularly FAA, suggest that cortical resting-state characteristics may also be relevant, but they were less robust and should be considered secondary. Overall, the results are consistent with the view that these sensory-affective experiences may be shaped not only by the characteristics of the stimulus, but also by the physiological state in which the stimulus is encountered.

## Notes

### Competing Interest Statement

The authors have declared no competing interest.

